# Raspberry Pi Powered Imaging for Plant Phenotyping

**DOI:** 10.1101/183822

**Authors:** Jose C. Tovar, J. Steen Hoyer, Andy Lin, Allison Tielking, Monica Tessman, Michael Miller, Steven T. Callen, S. Elizabeth Castillo, Noah Fahlgren, James C. Carrington, Dmitri A. Nusinow, Malia A. Gehan

## Abstract

- *Premise of the study*: Image-based phenomics is a powerful approach to capture and quantify plant diversity. However, commercial platforms that make consistent image acquisition easy are often cost-prohibitive. To make high-throughput phenotyping methods more accessible, low-cost microcomputers and cameras can be used to acquire plant image data.
- *Methods and Results*: We used low-cost Raspberry Pi computers and cameras to manage and capture plant image data. Detailed here are three different applications of Raspberry Pi controlled imaging platforms for seed and shoot imaging. Images obtained from each platform were suitable for extracting quantifiable plant traits (shape, area, height, color) *en masse* using open-source image processing software such as PlantCV.
- *Conclusion*: This protocol describes three low-cost platforms for image acquisition that are useful for quantifying plant diversity. When coupled with open-source image processing tools, these imaging platforms provide viable low-cost solutions for incorporating high-throughput phenomics into a wide range of research programs.

## INTRODUCTION

Image-based high-throughput phenotyping has been heralded as a solution for measuring diverse traits across the tree of plant life (Araus and Cairns, 2014; Goggin et al., 2015). In general, there are five steps in image-based plant phenotyping: 1) image and metadata acquisition; 2) data transfer; 3) image segmentation (separation of target object and background); 4) trait extraction (object description); and 5) population-level data analysis. Image segmentation, trait extraction, and data analysis are the most time-consuming steps of the phenotyping process, but protocols that increase the speed and consistency of image and metadata acquisition greatly speed up downstream analysis steps. Commercial high-throughput phenotyping platforms are powerful tools to collect consistent image data and metadata and are even more effective when designed for targeted biological questions (Topp et al., 2013; Chen et al., 2014; Honsdorf et al., 2014; Yang et al., 2014; Al-Tamimi et al., 2016; Pauli et al., 2016; Feldman et al., 2017; Zhang et al., 2017). However, commercial phenotyping platforms are cost-prohibitive to many laboratories and institutions. There is also no such thing as a ‘one-size fits all’ phenotyping system; different biological questions often require different hardware configurations. Therefore, low-cost technologies that can be used and repurposed for a variety of phenotyping applications are of great value to the plant community.

Raspberry Pi computers are small (credit card sized or smaller), low-cost, and were built as educational tools (Upton and Halfacree, 2014). Several generations of Raspberry Pi single-board computers have been released, and most models now feature built-in modules for wireless and bluetooth connectivity (Monk, 2016). The Raspberry Pi foundation also releases open source software and accessories such as camera modules (5 and 8 megapixel). Additional sensors or controllers can be connected via USB ports and general-purpose input/output pins. A strong online community of educators and hobbyists provide support (including project ideas and documentation), and a growing population of researchers use Raspberry Pi computers for a wide range of applications including phenotyping. We and others (e.g. Huang et al., 2016; Mutka et al., 2016; Minervini et al., 2017) have utilized Raspberry Pi computers in a number of configurations to streamline collection of image data and metadata. Here, we document three different methods for using Raspberry Pi computers for plant phenotyping (Figure 1.). These protocols are a valuable resource because while there are many phenotyping papers that outline phenotyping systems in detail (Granier et al., 2006; Iyer-Pascuzzi et al., 2010; Jahnke et al., 2016; Shafiekhani et al., 2017), there are few protocols that provide step-by-step instructions for building them (Bodner et al., 2017; Minervini et al., 2017). We provide examples illustrating automation of photo capture with open source tools (based on Python and standard Linux utilities). Further, to demonstrate that that these data are of high quality and suitable for quantitative trait extraction, we segmented example image data (plant isolated from background) using the open-source open-development phenotyping software PlantCV (Fahlgren et al., 2015).

**Figure 1.**
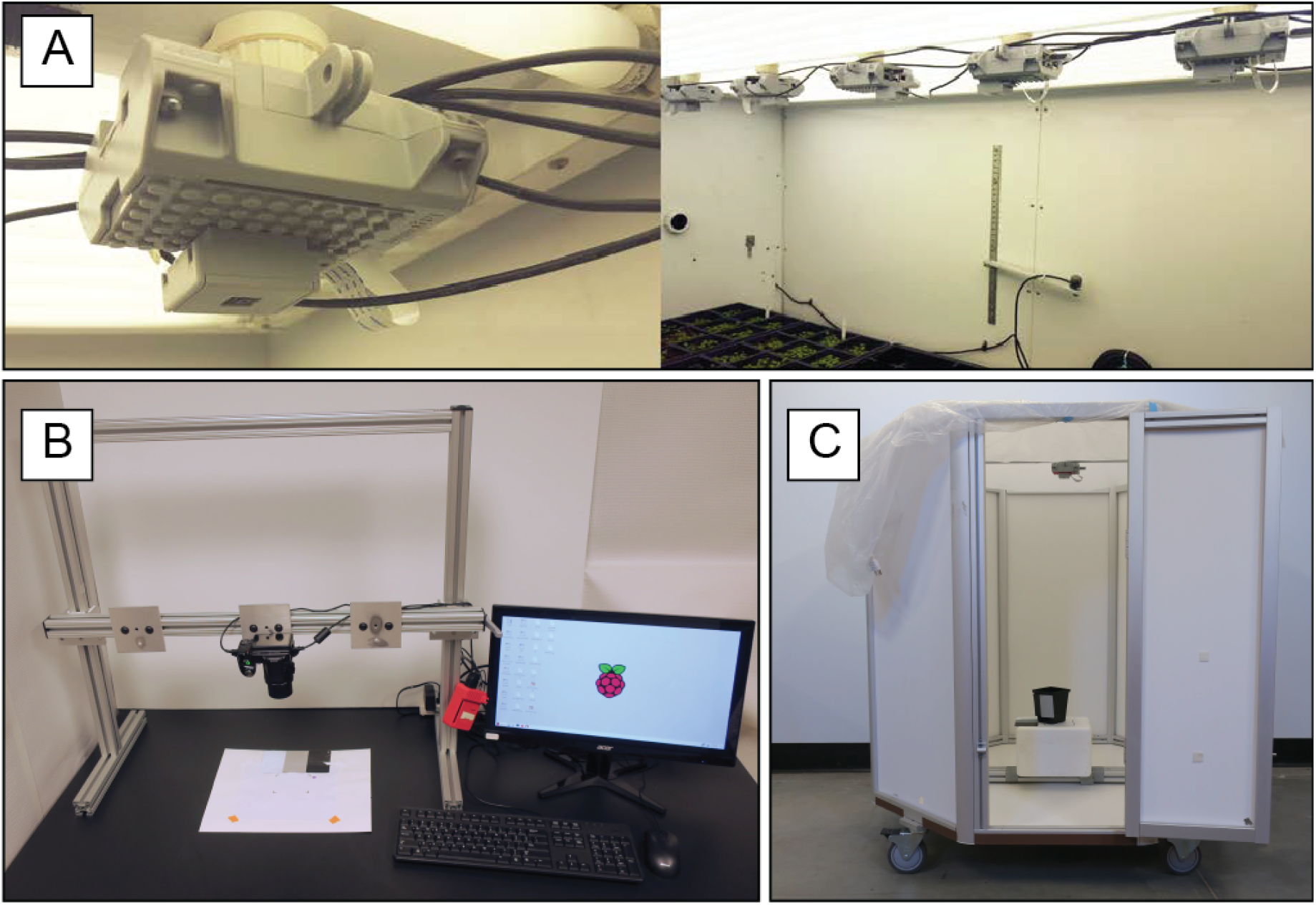
Low-cost Raspberry Pi phenotyping platforms. A) Raspberry Pi time-lapse imaging in a growth chamber. B) Raspberry Pi camera stand. C) Raspberry Pi multi-image octagon.

## METHODS AND RESULTS

### Raspberry Pi Initialization

This work describes three protocols (Appendices 2–4) that utilize Raspberry Pi computers for low-cost image-based phenotyping and gives examples of the data they produce. Raspberry Pi computers can be reconfigured for different phenotyping projects and can be easily purchased from online retailers. The first application is time-lapse plant imaging (Appendix 2); the second protocol describes setup and use of an adjustable camera stand for top-view photography (Appendix 3); and the third project describes construction and use of an octagonal box for acquiring plant images from several angles simultaneously (Appendix 4). For all three phenotyping protocols, the same protocol to initialize Raspberry Pi computers is used and is provided in Appendix 1. The initialization protocol in Appendix 1 parallels the Raspberry Pi Foundation’s online documentation and provides additional information on setting up passwordless secure shell (SSH) login to a remote host for data transfer and/or to control multiple Raspberry Pis. Passwordless SSH allows one to pull data from the data collection computer to a remote server without having to manually enter login information each time. Reliable data transfer is an important consideration in plant phenotyping projects because, while it is possible to process image data directly on a Raspberry Pi computer, most users will prefer to process large image datasets on a bioinformatics cluster. Remote data transfer is especially important for time-lapse imaging setups, such as the configuration described in Appendix 2, because data can be generated at high frequency over the course of long experiments, and thus can easily exceed available disk space on the micro secure digital (SD) cards that serve as local hard-drives. Once one Raspberry Pi has been properly configured and tested, the fully configured operating system can be backed up, yielding a disk image that can be copied (“cloned”) onto as many additional SD cards as are needed for a given phenotyping project (Appendix 1).

### Raspberry Pi Time-lapse Imaging

Time-lapse imaging is a valuable tool for documenting plant development and can reveal differences that would not be apparent from endpoint analysis. Raspberry Pi computers and camera modules work effectively as phenotyping systems in controlled-environment growth chambers; and low cost of Raspberry Pi computers allows this approach to scale well. Growth chambers differ from (agro)ecological settings but are an essential tool for precise control and reproducible experimentation (Poorter et al., 2016). Time-lapse imaging with multiple cameras allows for simultaneous imaging of many plants and can capture higher temporal resolution than conveyor belt and mobile-camera systems. Appendix 2 provides an example protocol for setting up the hardware and software necessary to capture plant images in a growth chamber. The main top-view imaging setup described is aimed at imaging flats or pots of plants in a growth chamber. We include instructions for adjusting the camera-plant focal distance (yielding higher plant spatial resolution) and describe how to adjust the temporal resolution of imaging. The focal distance can be optimized to the target plant, trait, and degree of precision required; large plant-camera distances allow a larger field of view, at the cost of lower resolution. For traits like plant area, where segmentation of individual plant organs is not critical, adjusting the focal length might not be necessary. Projected leaf area in top-down photos correlates well with fresh and dry weight, especially for relatively flat plants such as *Arabidopsis thaliana* (Leister et al., 1999). A stable and level imaging configuration is important for consistent imaging across long experiments and to compare data from multiple Raspberry Pi/Camera ris. Although there is more than one way to suspend Raspberry Pi/Camera rigs in a flat and stable top-view configuration, AC power socket adapters were attached to the the back of cases with silicone adhesive (Appendix 2). Raspberry Pi boards and cameras were then encased and screwed into the incandescent bulb sockets built into the growth chamber (Figure 1). Users with access to a 3D printer may prefer to print cases, so we have provided a link to instructions for printing a suitable case (with adjustable ball-joint Raspberry Pi camera module mount) in Appendix 2. This type of 3D printed case also works well for side-view imaging of plants grown on plates (Huang et al., 2016; Mutka et al., 2016). For this top-down imaging example, twelve Raspberry Pi/Camera rigs were powered through two USB power supplies drawing power (via extension cord and surge protector) from an auxiliary power outlet built into the growth chamber. Time-lapse imaging was scheduled at five-minute intervals using the software utility cron. A predictable file naming scheme that includes image metadata (field of view number, timestamp, and a common identifier) was employed to confirm that all photo timepoints were captured and transferred as scheduled. Images were pulled from each Raspberry Pi to a remote server twice per hour (using a standard utility called rsync) by a server-side cron process using the configuration files described in Appendix 2.

Optimizing imaging conditions for maximum consistency can simplify downstream image processing. To aid in image normalization during processing, color standards and size markers can be included in images. Placing rubberized blue mesh (Con-Tact Brand, Pomona, California, USA) around the base of plants can sometimes simplify segmentation (i.e. distinguishing plant foreground pixels from soil background pixels), though this was not necessary for the *A. thaliana* example described here. Care should be taken to ensure that large changes in the scene (including gradual occlusion of blue mesh by leaves) do not dramatically alter automatic exposure and color balance settings over the course of an experiment. If automatic exposure becomes an issue, camera settings can be manually set (see Appendix 4). In this example, cameras and flats were set up to yield a similar vantage point (a 4 x 5 grid of pots) in each field of view, such that very similar computational pipelines can be used to process images from all twelve cameras. An example image has been processed with PlantCV (Fahlgren et al., 2015), and the analysis script is available at https://github.com/danforthcenter/apps-phenotyping (Figure 2).

**Figure 2.**
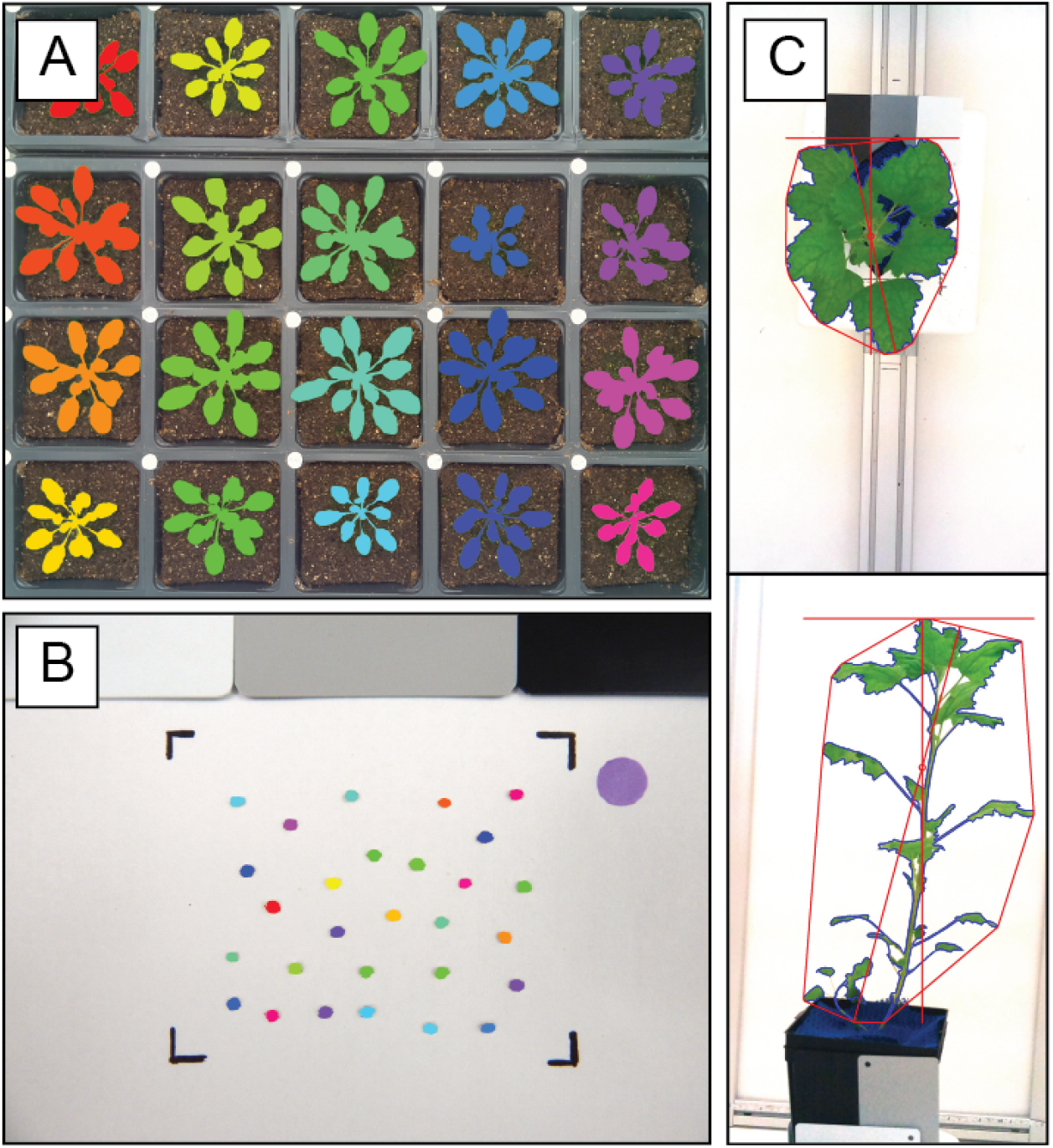
Examples of data collected from Raspberry Pi phenotyping platforms that have plant/seed tissue segmented using open-source open-development software PlantCV (Fahlgren et al., 2015). A) PlantCV-segmented image of a flat of Arabidopsis acquired from Raspberry Pi time-lapse imaging protocol in a growth chamber. B) PlantCV-segmented image of quinoa seeds acquired from Raspberry Pi camera stand. C) Example side- and top-view images of quinoa plants acquired from Raspberry Pi multi-image octagon. Plant convex hull, width, and length, have been identified with PlantCV and are denoted in red.

### Raspberry Pi Camera Stand

An adjustable camera stand is a versatile piece of laboratory equipment for consistent imaging. Appendix 3 is a protocol for pairing a low-cost home-built camera stand with a Raspberry Pi computer for data capture and management. The camera stand (79 cm width x 82.5 cm height) was built from aluminum framing (80/20, Columbia City, Indiana, USA) to hold a Nikon Coolpix L830 camera via a standard mount (Figure 1). For this application, we prefer to use a single-lens reflex (SLR) digital camera (rather than a Raspberry Pi camera module) for adjustable focus and to improve resolution. The camera was affixed to a movable bar, so the distance between camera and object can be adjusted up to 63 cm. A Python script that utilizes gphoto2 (Figuière and Niedermann, 2017) for data capture and rsync for data transfer to a remote host is included in the protocol (Appendix 3). When the ‘camerastand.py’ script is run, the user is prompted to enter the filename for the image. The script verifies that the camera is connected to the Raspberry Pi, acquires the image with the SLR camera, retrieves the image from the camera, renames the image file to the user-provided filename, saves a copy in a local Raspberry Pi directory, and transfers this copy to the desired directory on a remote host. As image filenames are commonly used as the primary identifier for downstream image processing, it is advised to use a filename that identifies the species, accession, treatment, and replicate, as appropriate. The Python script provided appends a timestamp to the filename automatically. We regularly use this Raspberry Pi Camera Stand to image seeds, plant organs (e.g. inflorescences), and short-statured plants. For seed images, a white background with a demarcated black rectangular area ensures that separated seeds are in frame, which speeds up the imaging process. Color cards (white, black, and gray; DGK Color Tools, New York, New York, USA) and a size marker to normalize area are also included in images to aid in downstream processing and analysis steps. It is advised to use the same background, and, if possible, the same distance between object and camera for all images in an experimental set. However, including a size marker in images can be used to normalize data extracted from images if the vantage point does change. *Chenopodium quinoa* (quinoa) seed images are shown as example data from the camera stand (Figure 2). Seed images acquired with the camera stand were processed using PlantCV (Fahlgren et al., 2015) to quantify individual seed size, shape, color, and count; these types of measurements are valuable for quantifying variation within a population. This overall process (Appendix 3) provides a considerable cost savings relative to paying for seed imaging services or buying a commercial seed imaging station.

### Raspberry Pi Multi-Image Octagon

Different plant architecture types require different imaging configurations for capture. For example, top-down photographs can capture most of the information about the architecture of rosette plants (as described above), but plants with orthotropic growth such as rice or quinoa are better captured with a combination of both side-view and top-view images. Therefore, platforms for simultaneously imaging plants from multiple angles are valuable. In Appendix 4, a protocol is described to set up an octagon-shaped chamber for imaging at different angles. A ‘master’ Raspberry Pi computer with a Raspberry Pi camera module is used collect image data and also to trigger three other Raspberry Pi computers/cameras. Data is transferred from the four Raspberry Pi computers to a remote host using rsync. The octagon chamber (122 cm height and 53.5 cm of each octagonal side) was constructed from aluminum framing and 3mm white pvc panels (80/20, Columbia City, Indiana, USA; Figure 1). The top of this structure is left open but is covered with a translucent white plastic tarp to diffuse light when acquiring images. A latched door was built into the octagon chamber to facilitate loading of plants. Four wheels were attached at the bottom of the chamber for mobility. The four Raspberry Pis with Raspberry Pi camera modules (one top-view and three side-views approximately 45° angle apart) in cases (SmartiPi Raspberry Pi B+ and camera case, Smarticase LLC, Philadelphia, Pennsylvania, USA) were affixed to the octagon chamber using heavy-duty velcro. To maintain a consistent distance between the Raspberry Pi cameras and a plant within the Raspberry Pi multi-image octagon, a pot was affixed to the center of the octagon chamber, with color cards affixed to the outside of the stationary pot (white, black, and gray; DGK Color Tools, New York, New York, USA) so that a potted plant could be quickly placed in the pot during imaging.

To facilitate data acquisition and transfer on all four Raspberry Pis, scripts are written so the user only needs to interact with a single ‘master’ Raspberry Pi (here the master Raspberry Pi is named ‘octagon’). From a laptop computer one would connect to the ‘master’ pi via SSH, then run the ‘sshScript.sh’ on that Pi. The ‘sshScript.sh’ script triggers the image capture and data transfer sequence in all four Raspberry Pis and appends the date to a user inputted barcode. When the ‘sshScript.sh’ script is run, a prompt asks the user for a barcode sequence. The barcode can be inputted manually, or, if a barcode scanner is available (we use a Socket 7Qi barcode scanner), a barcode scanner can be used to input the filename information. Again, it is advised to use a plant barcode that identifies the species, accession, treatment, and replicate, as appropriate. Once a barcode name has been inputted, another prompt asks if the user would like to continue with image capture. This pause in the ‘sshScript.sh’ script gives the user the opportunity to place the plant in the octagon before image capture is triggered. The sshScript.sh runs the script piPicture.py on all four Raspberry Pis. The ‘piPicture.py’ script captures an image and appends the user inputted filename with the Raspberry Pi camera id and the date. The image is then saved to a local directory on the Raspberry Pi. The ‘syncPi.sh’ script is then run by ‘sshScript.sh’ to transfer the images from the four Raspberry Pis to a remote host. The final script (shutdown_all_pi) is optionally run when image acquisition is over, allowing the user to shut down all four Raspberry Pis simultaneously. Examples of quinoa plant images captured with the Raspberry Pi multi-image octagon are provided and analyzed with PlantCV (Fahlgren et al., 2015) for plant area and shape (Figure 2).

### Protocol Feasibility

The protocols provided in the appendices that follow provide step-by-step instructions for using Raspberry Pi computers for plant phenotyping in three different configurations. The majority of components for all three protocols are readily available online. Low-cost computers and components are especially important since some experiments might test harsh environmental conditions and need to be replaced long-term. Each of the platforms were built and programmed in large part by high-school students, undergraduates, or graduate students and do not require a large investment of time to build or set-up. Automation increases the consistency of image and metadata capture, which streamlines image segmentation (Figure 2) and is thus preferable to manual image capture. Furthermore, the low cost of each system and the flexibility to reconfigure Raspberry Pi computers for multiple purposes makes automated plant phenotyping accessible to most researchers.

## CONCLUSION

The low-cost imaging platforms presented here provide an opportunity for labs to introduce phenotyping equipment into their research toolkit, and thus increase the efficiency, reproducibility, and thoroughness of their measurements. These protocols make high-throughput phenotyping accessible to researchers unable to make a large investment in commercial phenotyping equipment. Paired with open-source open-development high-throughput plant phenotyping software like PlantCV (Fahlgren et al., 2015), image data collected from these phenotyping systems can be used to quantify plant traits for populations of plants that are amenable to genetic mapping. These Raspberry-Pi-powered tools are also useful for education and training. In particular, we have used time-lapse imaging to introduce students and teachers to the Linux environment, image processing, and data analysis in a classroom setting (http://github.com/danforthcenter/outreach/). As costs continue to drop and hardware continues to improve, there is enormous potential for the plant science community to capitalize on creative applications, well-documented designs, and shared datasets and code.

## Acknowledgements

This work was funded by the Danforth Plant Science Center and the National Science Foundation: IOS-1456796,10S-1202682, EPSCoR IIA-1355406, IIA-1430427, IIA-1430428, MCB-1330562, DBI-1156581. We thank Neff Power (St. Louis, MO) for help in determining the framing to build the multi-image octagon and camera stand. We thank George Wang (MPI Tübingen; Computomics Corporation), César Lizárraga, Leonardo Chavez, and Nadia Shakoor for helpful discussions and the editors of this series for the opportunity to contribute this paper.

## Appendix 1. Initializing a Raspberry Pi for phenotyping projects

The camera stand, growth-chamber imaging stations, and multi-image octagon phenotyping platforms that are described in detail in Appendices 2-4, use Raspberry Pis to trigger image acquisition, append metadata to filenames, and to move data. The following are the required parts and steps to initialize a single Raspberry Pi. The initialization protocol is based on the installation guidelines from the Raspberry Pi Foundation, which are under a Creative Commons license (https://www.raspberrypi.org/documentation/).

Parts List

*Parts List:*
| Item |
| --- |
| Raspberry Pi single-board microcomputer |
| Micro USB power supply |
| Mini Secure Digital (SD) card, we recommend 16GB |
| HDMI Monitor, HDMI cable, keyboard, and mouse (for initialization process) |

### General Raspberry Pi Initialization Protocol

1. Install ‘Raspbian Stretch with Desktop’ (here version 4.9 is used, but the latest version is recommended) onto the SD card by following the installation guide at https://www.raspberrypi.org/downloads/raspbian/
2. Insert mini SD card into Raspberry Pi and plug in monitor, keyboard, and mouse to Raspberry Pi.
3. Plug in Micro USB power supply and connect to power. The Raspberry Pi will boot to the desktop interface, which is also known as the graphical user interface (GUI). If the Raspberry Pi does not boot to the desktop interface, you can type sudo raspi-config and go to the third option ‘Enable Boot to Desktop/Scratch’ to change this. Alternately, if the Raspberry Pi boots to the command line you can get to the GUI by typing ‘startx’ and enter.
4. Once at the desktop, open Raspberry Pi Configuration under Applications Menu > Preferences. Alternatively, you can get to the configurations menu by typing “sudo raspi-config” in the Terminal program.

a. In the System tab, set hostname (see Appendices 2, 3, or 4 for specific hostnames to use; alternatively, a static IP address can be set up for the Raspberry PI).
b. In the Interfaces tab, set SSH and Camera to enabled.
c. In the Localization tab:

i. Set Locale to appropriate Language and Country, and leave Character Set as UTF-8 (default option).
ii. Set Timezone to appropriate Area and Location.
iii. Set Keyboard to appropriate Country and Variant.
iv. Set WiFi Country.
5. Configure WiFi using the network icon on the top right of the desktop. Alternatively, use an Ethernet cable connection.
6. Optionally, make a local copy of the scripts that accompany this paper. In Terminal, change directory to the Desktop by typing “cd Desktop”. Then type “git clone https://github.com/danforthcenter/apps-phenotyping.git”. If prompted with “The authenticity of host ‘remote-hosť can’t be established (…) Are you sure you want to continue connecting?” enter “yes”. This will download the project scripts and examples for all three phenotyping platforms (Appendices 2-4). Some of these scripts may need be adjusted after they have been cloned on the Raspberry Pi, as described below.

### Raspberry Pi SD card cloning protocol

Once you have gone through the initialization protocol for one Raspberry Pi the disk image of the SD card from that Raspberry Pi can be cloned if you need additional Raspberry Pis for your project. Any project specific scripts that need further adjustments on individual Raspberry Pis can then be completed (see Appendices 2 to 4). Cloning an SD card will generate a file of the exact size of the SD card (e.g. 16 GB). Make sure the SD cards to which the original image will be cloned are at least the same size as the original initialized card (e.g. 16 GB or bigger).

### To clone an SD card on a Windows computer

1. Download and install Win32 Disk Imager from https://sourceforge.net/projects/win32diskiiriager/
2. Before opening the Win32 Disk Imager, insert the SD card (in an SD card reader if needed) from the initialized Raspberry Pi into your computer.
3. Open Win32 Disk Imager.
4. Click on the blue folder icon. A file explorer window will appear.
5. Select the directory to store the SD card image, and provide a filename for the image.
6. Click Open to confirm your selection. The file explorer window will close.
7. Under Device, select the appropriate drive letter for the SD card.
8. Click the Read button.
9. Once the image is created, a ‘Read Successful’ message will appear. Click OK.
10. Eject the SD card, and close Win32 Disk Imager.
11. Insert the new SD card where the image will be cloned. Make sure this SD card has as much or more storage capacity as the SD card from the initialized Raspberry Pi that was imaged.
12. Reopen Win32 Disk Imager.
13. Click on the blue folder icon, and select the image that was just created.
14. Under Device, select the appropriate drive letter for the SD card where the image will be cloned.
15. Click the Write button.
16. Click Yes.
17. Once the image is created, a ‘Write Successful’ message will appear. Click OK.
18. Eject the SD card, and insert it into the Raspberry Pi. The Raspberry Pi is now initialized.

### To clone an SD card on a Mac computer

1. Download and install ApplePi-Baker from https://www.tweaking4all.com/software/macosx-software/macosx-apple-pi-baker/
2. Insert the SD card (in an SD card reader if needed) from the initialized Raspberry Pi into your computer.
3. Under Pi-Crust: Select SD-Card or USB drive, select the initialized Raspberry Pi SD card.
4. Click on Create Backup.
5. Click OK.
6. Under Save As, provide a filename for the SD card image.
7. Under Where, select directory to store the SD card image, and click Save.
8. Once the image is created, a ‘Your ApplePi is Frozen! ‘ message will appear. Click OK.
9. Eject the SD card.
10. Insert the new SD card where the image will be cloned. Make sure this SD card has as much or more storage capacity as the SD card from the initialized Raspberry Pi that was imaged.
11. Under Pi-Crust: Select SD-Card or USB drive, select the SD card where the image will be cloned.
12. Click on Restore Backup.
13. Browse and select the image that was just created.
14. Click OK.
15. Once the SD card is cloned, a ‘Your ApplePi is ready!’ message will appear.
16. Click OK.
17. Eject the SD card, and insert it into the Raspberry Pi. The Raspberry Pi is now initialized.

### To clone an SD card on a Linux computer

This protocol is adapted from The PiHut (https://thepihut.com/blogs/raspbeny-pi-tutorials/17789160-backing-up-and-restoring-your-raspberry-pis-sd-card) and Raspberry Pi Stack Exchange (https://raspberrypi.stackexchange.com/questions/311/how-do-i-backup-my-raspberry-pi)

1. First, use the command ‘df-h’ to see a list of existing devices.
2. Insert the SD card (in an SD card reader if needed) from the initialized Raspberry Pi into your computer.
3. Use the command ‘df-h’ again. The SD card will be the new item on the list (e.g. /dev/sdbp1 or /dev/sdb1). The last part of the name (e.g. p1 or 1) is the partition number.
4. Use the command ‘sudo dd if=/dev/SDCardName of=/path/to/SDCardlmage.img’ to create the SD card image (e.g. sudo dd if=/dev/sdb of=~/InitializedPi.img). Make sure to remove the partition name to image the entire SD card (e.g. use /dev/sdb instead of /dev/sdbl).
5. There is no progress indicator, so wait until the command prompt reappears.
6. Unmount the SD card by typing: sudo umount /dev/SDCardName.
7. Remove the SD card.
8. Insert the new SD card where the image will be cloned. Make sure this SD card has as much or more storage capacity as the SD card from the initialized Raspberry Pi that was imaged.
9. Use the command ‘df-h’ again to discover the new SD card name, or names if there is more than 1 partition.
10. Unmount every partition using the command ‘sudo umount /dev/SDCardName’ (e.g. sudo umount/dev/sdbl).
11. Copy the initialized Raspberry Pi SD card image using the command ‘sudo dd if=/path/to/SDCardImage.img of=/dev/SDCardName’.
12. There is no progress indicator, so wait until the command prompt reappears.
13. Unmount the SD card, and insert it into the Raspberry Pi. The Raspberry Pi is now initialized.

### General instructions for installing and testing a Raspberry Pi Camera

1. Make sure the Raspberry Pi is not connected to power.
2. Pull up the top part of the connector located between the HDMI and ethernet ports, until loose.
3. Insert the Raspberry Pi camera flex cable into the connector, with the silver rectangular plates at the end of the cable facing the HDMI port.
4. While holding the cable in place, push down the top part of the connector to prevent the flex cable from moving.
5. Remove the small piece of blue plastic covering the camera lens, if present.
6. Turn the Raspberry Pi on, and test the camera by opening a Terminal window, then entering “raspistill-o image-name-here.jpg” to take a picture.

### General instructions for using a SSH keys for passwordless connection to a remote host

These instructions are to allow a Raspberry Pi computer to access a remote host (e.g. another Raspberry Pi computer, bioinformatics cluster, or other computer), without having to enter login information. If a user would like to pull data from Raspberry Pi to a remote host, rather than pushing data from a Raspberry Pi to a remote host, similar instructions would be followed on the remote computer.

1. In the Terminal window, enter “ssh-keygen” to create a public SSH key for passwordless access to a remote host.
2. Press enter to use the default location when asked to “Enter file in which to save the key”.
3. Press enter two times, to use no passphrase.
4. Optionally, enter “Ls ~/.ssh” to verify the SSH key was generated. The files “id_rsa” and “id_rsa.pub” should be listed.
5. Use the command “ssh-copy-id-i ~/.ssh/id_rsa.pub user@remote-host” in the Terminal window to copy the public SSH key to the remote host, where “user@remote-host” should be replaced by the name of the remote host where the images will be stored (e.g. ssh-copy-id-i ~/.ssh/id_rsa.pub jdoe@serverx).
6. If prompted with “The authenticity of host ‘remote-hosť can’t be established (…) Are you sure you want to continue connecting?” enter “yes”.
7. Enter the user’s password for the remote server, if prompted.
8. Verify the SSH key was successfully copied to the remote host. SSH to the remote host from the Raspberry Pi, using the command “ssh user@remote-host” in the Terminal window (e.g. ssh jdoe@serverx). No password should be required if the key was copied successfully.

### Additional Notes

- For official distributions of the Raspbian Raspberry Pi operating system, the default username is “pi”, and default password is “raspberry”. The following protocols assume these default settings are unchanged. To change passwords, open Raspberry Pi Configuration under Applications Menu > Preferences. In the System tab, click on ‘Change Password…’. Enter current password, new password, and confirm new password. If using the command line type passwd, then follow the command prompts to change password.
- If an Ethernet cable connection is used, a Power over Ethernet adapter (such as UCTronics LS-POE-B0525) and a Power over Ethernet-capable Ethernet switch (such as Ubiquiti ES-48-750W) can be used, eliminating the need for Raspberry Pi power cables.

## Appendix 2. Raspberry Pi Top-View or Side-View Time-Lapse Imaging

The protocol that follows describes how to set up one or more Raspberry Pi/Camera rigs for time-lapse photography, and is based on the tutorial at https://www.raspberrypi.org/documentation/usage/camera/raspicam/timelapse.md

Here, we focus on a 12-camera configuration that has worked well in a reach-in growth chamber. We describe one low-cost method for stably fixing Raspberry Pi/Camera rigs to the top of the chamber. Zip ties may work well for attaching Pi/Camera rigs in some growth chambers ((Minervini et al., 2017), and we have also used heavy-duty velcro, so that Pi/Camera rigs can be removed and used for other purposes, such as side-view imaging of seedlings on petri plates. We have separately provided a protocol for imaging petri plates with a NoΓR camera module and IR light-emitting diode (LED) panel (as in Huang et al. 2016); see http://maker.danforthcenter.org/tutorial/raspberry%20pi/led/raspberry%20pi%20camera/RPi-LED-Illumination-and-Imaging

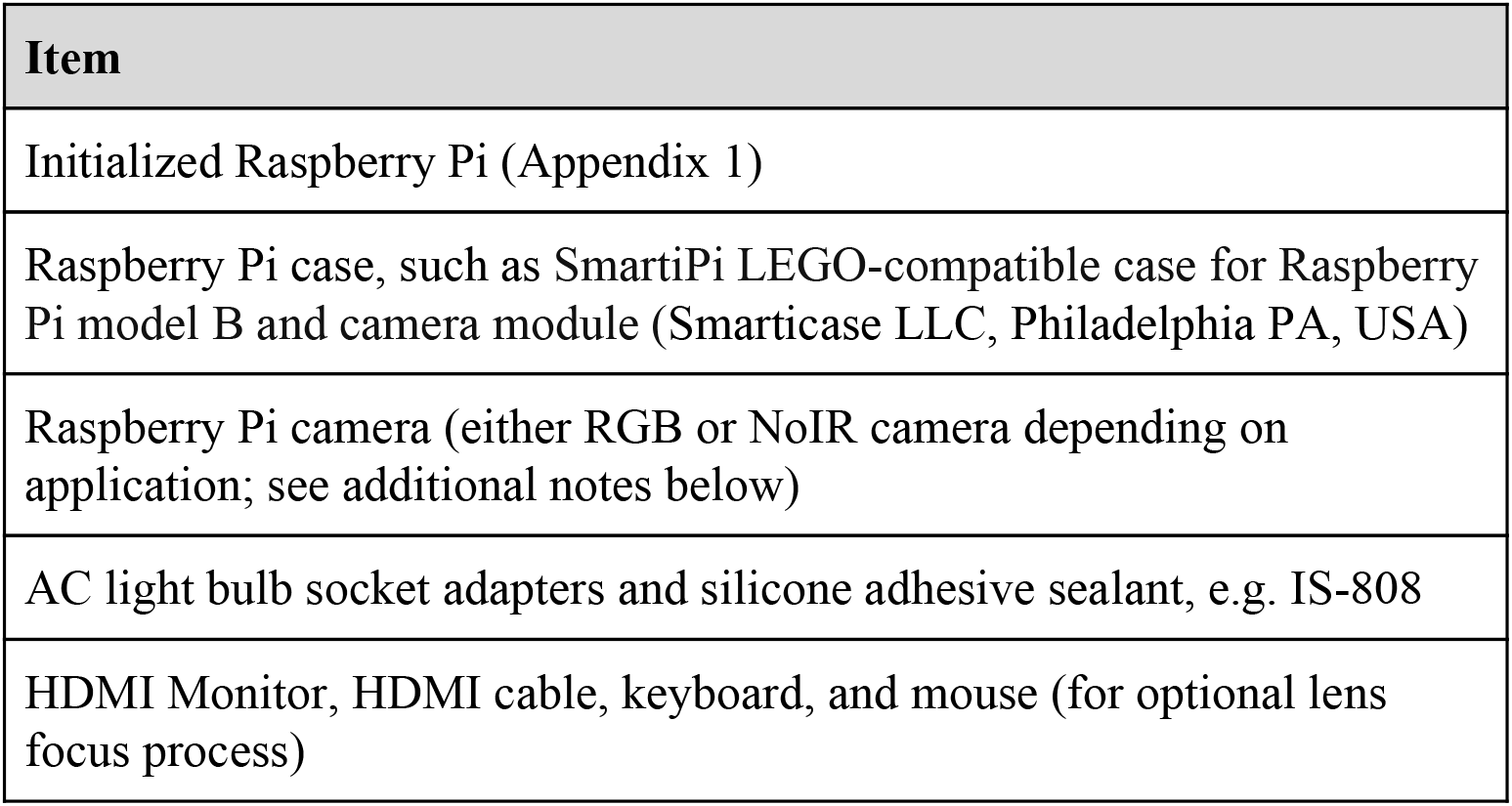
Parts list

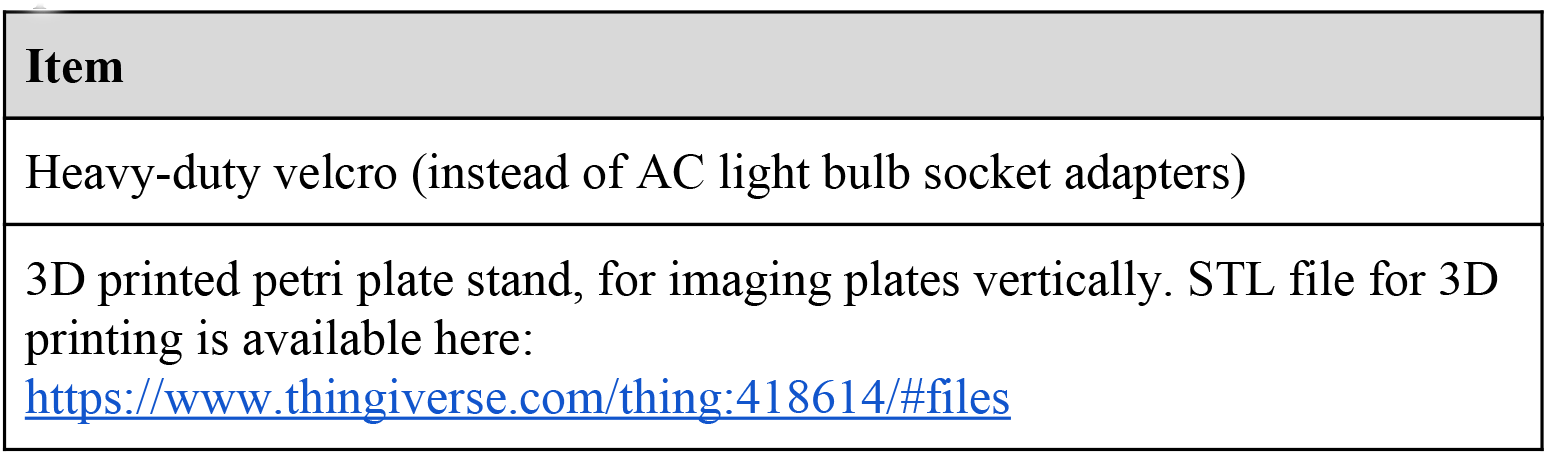
Optional/Alternative Parts

### Adjustment of Raspberry Pi Camera Focus (Optional)

As noted above, the Raspberry Pi camera is fixed focus, and therefore if you want to alter the focus you have to alter the Raspberry Pi camera module itself with pliers. We recommend watching the excellent youtube tutorial by George Wang: https://www.youtube.com/watch?v=u6VhRVH3Z6Y. For the main configuration described here, the lens-plant distance was 55.2 cm.

1. Install the Raspberry Pi camera on a Raspberry Pi computer that is plugged into a Monitor, Mouse and Keyboard as described above. Plug the power in last).
2. To change the focus, use a sharp object and carefully remove the glue around the lens. Every few incisions, carefully grip the lens with pliers and check if it will turn counter-clockwise. (Removing the glue is optional, as noted in the video linked above.)
3. When the camera lens does turn, use the following command in the Terminal of the Raspberry Pi to check whether a target is in focus at your desired imaging distance: “python raspistill- o image.jpg” We suggest using a ruler and white business card as targets. Continue adjusting the camera at the desired focal distance.

### Mounting of Raspberry Pi/Camera rigs for top-view time-lapse imaging

1. Connect a camera module to each Raspberry Pi, as described in Appendix 1.
2. Attach the AC light bulb socket adapters to the Raspberry Pi cases with silicone adhesive, and install an initialized Raspberry Pi computer (see Appendix 1) and camera in each case.
3. Plug each of the Raspberry Pis into the monitor, keyboard, mouse, and USB power. Following Appendix 1, change the hostnames of the Raspberry Pis. For example, the sample configuration files we provide assume that the twelve Raspberry Pi have hostnames ‘timepi0ľ, ‘timepi02, … ‘timepil2\ Recall that protocol 1 describes how to copy these configuration files, e.g. to /home/pi/Desktop.
4. Make a folder for images on each of the Raspberry Pis. In our example we made a folder ‘/home/pi/images’. To do this on the Terminal type: ‘mkdir /home/pi/images’. If you wish to use a different path image folder, change line 14 in the example file ‘pull-images-from-raspi.crontab’ (within the appendix.2.time-lapse subdirectory). You will also need to change the ‘photograph-all-5min.crontab’ and ‘photograph-all-5min-vhflipped.crontab’ lines 12, 15, 18, 21, and 24 so that the images are saved to the correct location during acquisition (both of these scripts are located in Desktop/apps-phenotyping/appendix.2.time-lapse)
5. Physically position each Pi/Camera rig within an experimental growth space (e.g. by screwing adapters into sockets or joining velcro strips together). Take photos (with the `raspistill` command) to confirm that each camera covers a suitable field of view. See additional notes below on taking photos remotely and optionally flipping photo orientation. ***Starting and ending a single imaging experiment:***
6. The ‘photograph-all-5min.crontab’ and ‘photograph-all-5min-vhflipped.crontab’ files are cron tables and contain the commands that trigger regular image capture. Both scripts are currently written to capture data every 5 minutes between the hours of 8:30 and 17:30 (8:30am to 5:30pm standard time). If that frequency is too high, the first number or comma separated list of numbers on lines 12, 15, 18, 21, and 24 have to be altered to reflect that change. If the hours of imaging are different, then the second number or range of numbers on lines 12, 15, 18, 21, and 24 has to be altered.
7. The ‘photograph-all-5min.crontab’ and ‘photograph-all-5min-vhflipped.crontab’ files also control wifi, turning off outside of the imaging window/photoperiod (see comments in files). Wifi is set to turn on ten minutes before the start of imaging and turn off 10 minutes after imaging ends. If the minute or hour of imaging is different from our experimental setup then the first two numbers on both lines 37 and 43 have to be altered.
8. Once both ‘photograph-all-5min.crontab’ and ‘photograph-all-5min-vhflipped.crontab’ scripts are satisfactory, install the cron jobs on each Raspberry Pi. The ‘install-twelve-crontabs.sh’ script (run from a remote machine, and depending on reasonable wifi connectivity) does this for all twelve Raspberry Pis, but first the user has to determine if the images need to be flipped or not. The difference between the ‘photograph-all-5min.crontab’ and ‘photograph-all-5min-vhflipped.crontab’ scripts is that the ‘photograph-all-5min-vhflipped.crontab’ imaging command flips the images in both the vertical and horizontal directions. Flipping the images might be necessary if there are differences in the orientation of the cameras, and thus images, between the Raspberry Pis. If a Raspberry Pi’s images are in ‘wrong’ orientation, open the ‘install-twelve-crontabs.sh’ file and follow the directions for commenting and uncommenting.
9. To pull data from the Raspberry Pi computers to a remote host, line 15 of the ‘pull-images-from-raspi.crontab’ must be changed to the path that you would like the images to go to on the remote host. The remote host must have passwordless SSH set up so that it can login to the Raspberry Pis without a password. This is very much like the ‘general instructions to generate a SSH key for passwordless SSH to a remote host’ in Appendix 1, but in reverse. Briefly, on the remote host you would generate an ssh key (command “ssh-keygen”), then copy that to the Raspberry Pi (e.g. “ssh-copy-id-i ~/.ssh/id_rsa.pub pi@timepi01”).
10. Once the RASPIDIR and SERVERDIR paths are changed in ‘pull-images-from-raspi.crontab’, put the “pull-images-from-raspi.crontab’ file on the remote host, then install it on the remote host on the command line by typing: ‘crontab pull-images-from-raspi.crontab’. Warning: this will overwrite existing cron jobs.
11. Upon conclusion of an experiment, suspend photography on each Raspberry Pi by “removing” the active crontab (crontab -r). Once the experiment is done, you can safely shutdown the Raspberry Pis using the ‘shut-down-all.sh’ script, if desired.
12. If a cron table job is set up on the Raspberry Pi it will take images as long as the Raspberry Pi has power, disk space, and a functioning camera module.

### Additional Notes

- Here we set up twelve Raspberry Pis for time-lapse imaging, but you may want to set up more or fewer. If fewer than twelve Raspberry Pis are used then simply comment out the excessive commands with a ‘#’ in each of the Appendix 2 scripts. If more than twelve Raspberry Pis are used, then follow the commented code to add more Raspberry Pis with unique hostnames.
- Because of the low cost of the hardware, this approach scales well. If you intend to use a large number of Raspberry Pis (tens, hundreds) you will likely want to investigate management and monitoring tools such as Ansible, Ganglia, and/or Puppet.
- Consider including at least one size marker in each field of view. For example, a white Tough-Spot (Research Products International, Mount Prospect, Illinois, USA) will remain affixed if wet.
- Also consider including color standards or white balance cards (white, gray and black; DGK Color Tools Optek Premium Reference White Balance Cards)
- As noted above, third generation Raspberry Pis have built in wifi and bluetooth modules. Older Rasbperry Pi models can be used with USB wifi modules (e.g. Newark 07W8938). Connectivity to our local wirelss network from within reach-in growth chambers is generally good, and has been more than sufficient for our monitoring and image transfer purposes. Testing wireless connectivity before setting up Pi/Camera rigs is strongly recommended. Wireless transfer is unlikely to work within a walk-in growth room, and with a large number of Pi/Camera rigs it is preferrable to avoid wireless signal interference by transferring data via ethernet. Alternatively, if real-time monitoring is not required and SD card disk space is not a constraint, one or more Raspberry Pis can be left to run autonomously until the conclusion of an experiment.
- If wireless connectivity is good, one can run test photo capture commands via a remote connection (and then copy the resulting image files for viewing, e.g. with rsync). This removes the need to physically connect a monitor and keyboard to check orientation when mounting each Pi/Camera rig. Minervini et al. ((Minervini et al., 2017) have provided instructions for installing and configuring an interface for taking and viewing photos through a web browser.
- The Raspberry Pi NoIR camera can be paired with an infrared (IR) light source to image under low visible light or no visible light conditions. We use a 730 nm cutoff filter (Lee #87) over the NoIR camera lens to block visible light when using an 880 nm LED array to backlight (see link above). The cutoff filter helps prevent changes in contrast during imaging, making image processing easier.

## Appendix 3. Raspberry Pi Camera Stand

The following are the hardware and software needed to set-up a Raspberry Pi camera stand.

Parts list

| Quantity | Description |
| --- | --- |
| 1 | Initialized Raspberry Pi Computer (see Appendix 1) |
| 1 | Raspberry Pi case |
| 1 | HDMI cable, mouse, keyboard, and monitor. |
| 1 | Nikon Coolpix L830 or other gphoto2 (Figuière and Niedermann, 2017) supported camera |
| 1 | Nikon Coolpix power cord |
| 1 | Nikon Coolpix L830 USB cable |

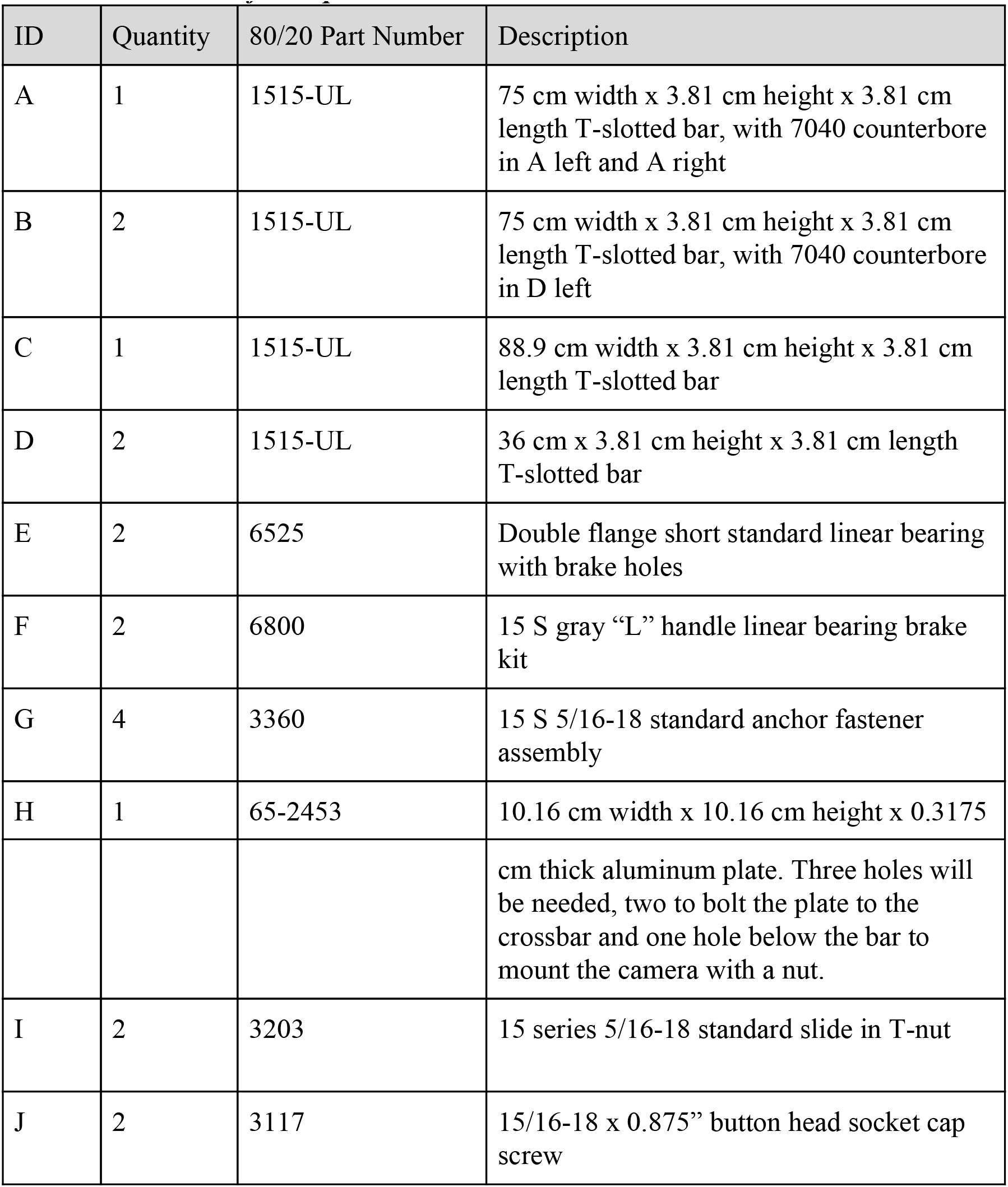
Aluminum 80/20 Inc. frame parts

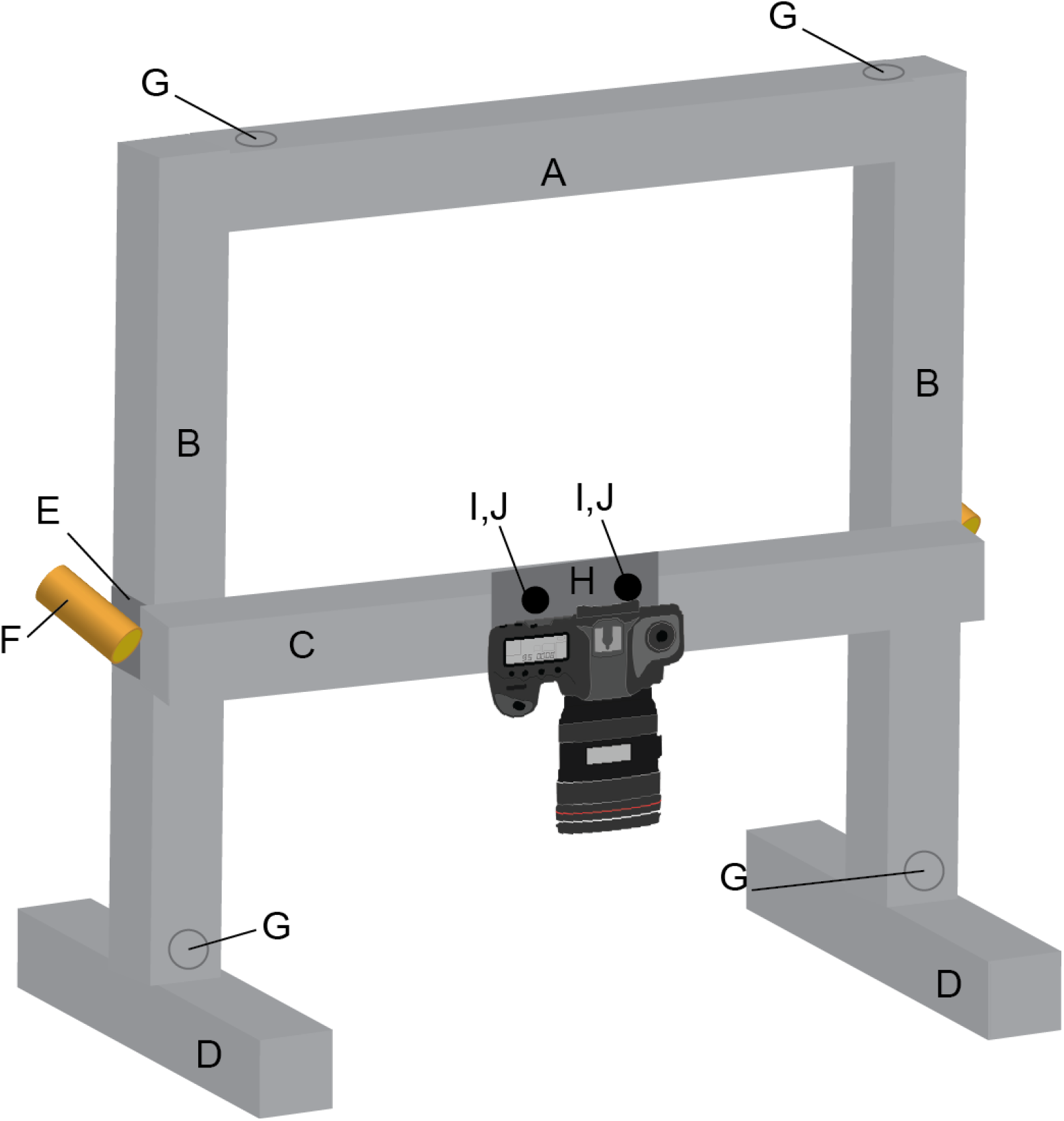

### Additional set-up of Raspberry Pi for camera stand

1. In the Applications Menu > Preferences, set hostname to camerastand.
2. Install gphoto2 and libgphoto2 (here stable version 2.5.14 is used, (Figuière and Niedermann, 2017)) by following the installation guide at https://github.com/gonzalo/gphoto2-updater.
3. Connect the camera to the Raspberry Pi with the Nikon Coolpix L830 USB cable.
4. Plug in and turn the camera on.
5. Open a Terminal window, enter “gphoto2-auto-detect” to detect the camera.
6. Optionally, in the Terminal window, enter “gphoto2-summary” to verify gphoto2 has correctly identified the Nikon Coolpix L830.
7. In the Terminal window, enter “cd Desktop” to change directory to the desktop.
8. Create a folder to store images on the Raspberry Pi desktop, and change the picPath directory in line 25 of camerastand.py to this folder (i.e. /home/pi/Desktop/folder1/)
9. Replace user@remote-host:remote-directory in line 32 of camerastand.py to the camera stand operator’s username, the remote host name, and the directory in the remote host where the images will be stored (e.g. jdoe@serverx:/home/jdoe/camerastand_images). Make sure that an SSH keygen has been generated (see Appendix 1) that will allow the Raspberry Pi to push data to the remote host.

### Raspberry Pi camera stand operation protocol

1. Turn the Raspberry Pi and camera on.
2. Open a Terminal window.
3. Change directory to Desktop (type “cd Desktop”).
4. In the Terminal, use the command “python /home/pi/Desktop/apps-phenotyping/appendix.3.camerastand/camerastand.py filename” then press enter to acquire and transfer an image, where “filename” should be replaced by an appropriate filename for the current picture (e.g. python /home/pi/Desktop/apps-phenotyping/appendix.3.camerastand/camerastand.py speciesx_plant 1 treatment 1 rep 1).

### Additional Notes

- The camera stand allows camera height to be adjusted. We recommend including a size marker in the images to normalize object area during image analysis. We often use a 1.27 cm diameter Tough-Spot (Research Products International, Mount Prospect, Illinois, USA).
- For seed image background, we draw the corners of a box on a white piece of paper or cardboard. We then place color cards (white, gray and black; DGK Color Tools Optek Premium Reference White Balance Card), and the size marker (Tough-Spot; Research Products International, Mount Prospect, Illinois, USA) just outside the box. This ensures that objects to be imaged (e.g. seeds) are within the field of view.
- Images are saved on the Raspberry Pi, as well as in the remote host, in the directories indicated in lines 25 and 32 of camerastand.py, respectively. Alternatively, the rsync command can be changed so that data is deleted from the Raspberry Pi once data transfer has been confirmed. To change the rsync command so that the image is deleted from the Raspberry Pi once it has been transferred to the remote host, change line 32 in the ‘camerastand.py’ script to “sp.call([“rsync”, “-uhrtP”, picPath, “user@remote-host:remote-directory”, remove-source-files”])”

## Appendix 4. Raspberry Pi Multi-Image Octagon

Parts list

*Parts list:*
| Quantity | Description |
| --- | --- |
| 1 | Laptop or other computer (could be a Raspberry Pi with a monitor, keyboard, and mouse) |
| 4 | Initialized Raspberry Pi computers (see Appendix 1) |
| 4 | Raspberry Pi cases (e.g. SmartiPi brand LEGO-compatible cases for Raspberry Pi model B case and camera module; SmartiCase LLC, Philadelphia PA, USA) |
| 4 | Raspberry Pi cameras (RGB) |
| 4 | Heavy-duty velcro |
| 1 | Power strip |
| 1 | White translucent tarp for light diffusion |
| 1 | HDMI Monitor, HDMI cable, keyboard, and mouse (for initialization process) |

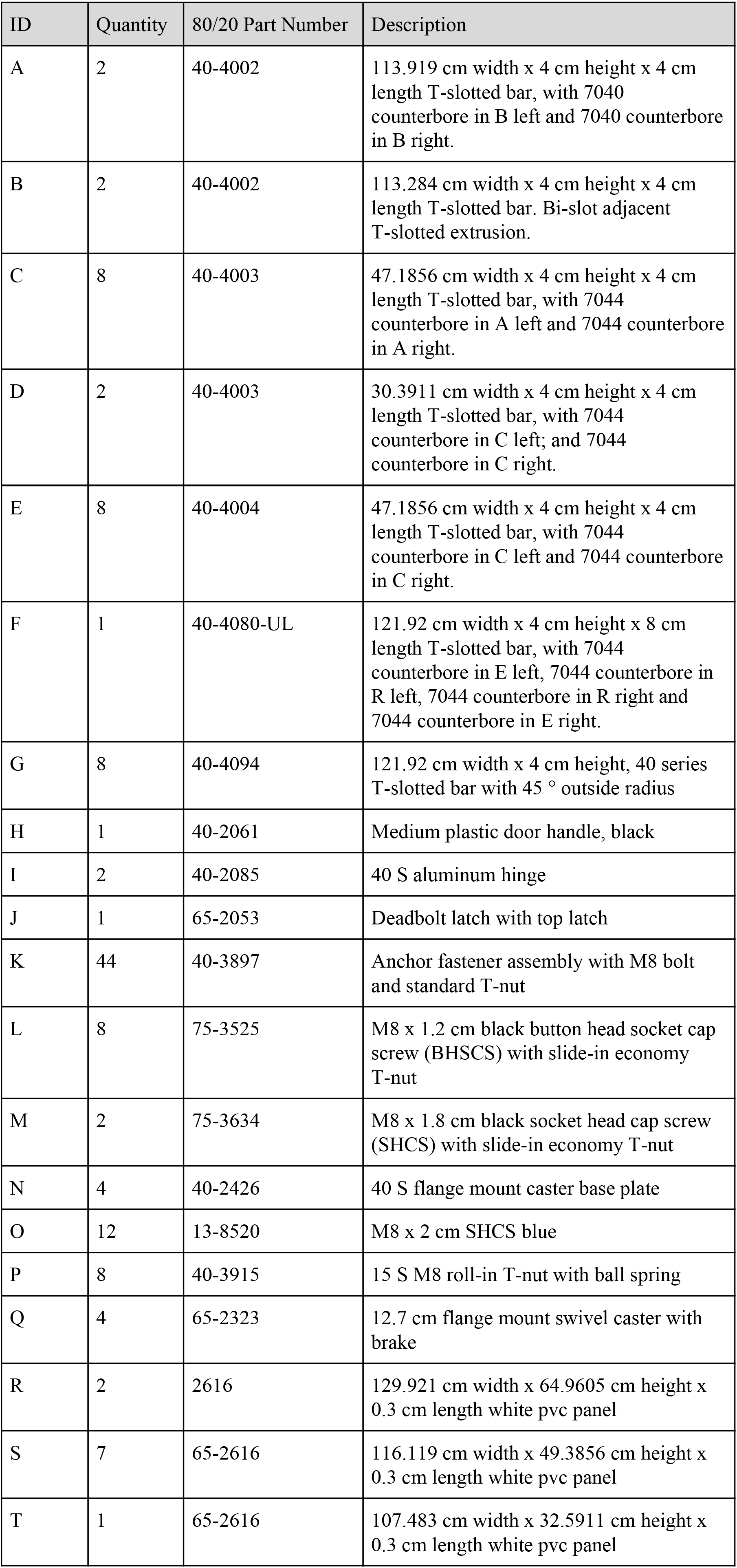
Aluminum 80/20 Inc. frame parts and paneling for Octagon

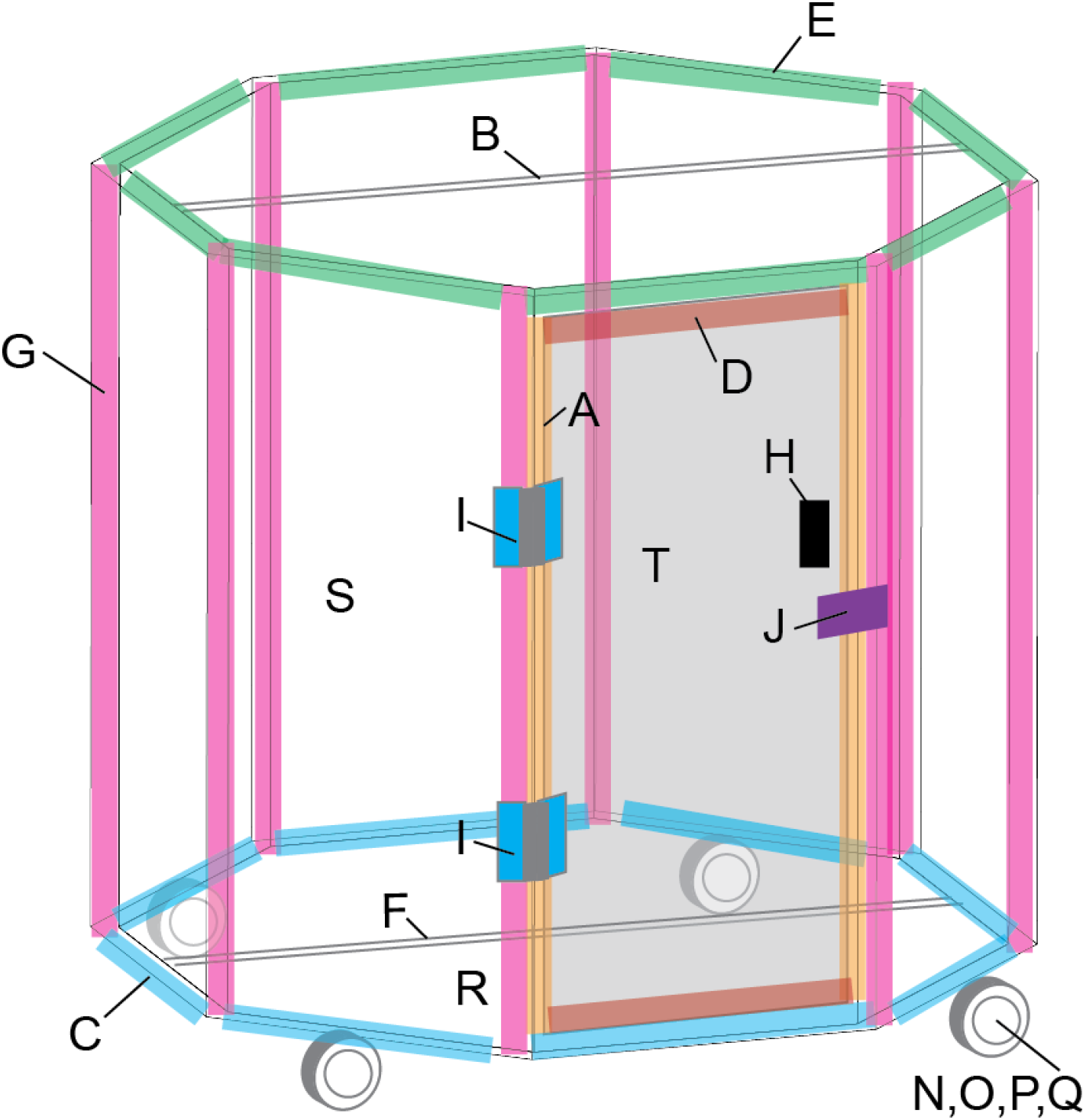

Optional parts

**Optional parts:**
| Quantity | Description |
| --- | --- |
| 1 | Barcode scanner (e.g. Socket Mobile 7Qi) |

### Additional set-up for 4 Raspberry Pis for multi-image octagon

1. Install Raspberry Pi camera on Raspberry Pi (see Appendix 1).
2. Plug each of the Raspberry Pis into the monitor, keyboard, mouse, and USB power. Following Appendix 1, set hostnames of the four Raspberry Pis to: 1) octagon; 2) sideview1; 3) sideview2; and 4) topviewpi
3. For the Raspberry Pi with the hostname ‘octagon’, set up passwordless SSH (Appendix 1) so that the ‘octagon’ Raspberry Pi can trigger scripts on other Raspberry Pis commands. Briefly:

a. open a Terminal window, and use the command “ssh-copy-id-i ~/.ssh/id_rsa.pub user@remote-host” for the three other Raspberry Pis (e.g ssh-copy-id-i ~/.ssh/id_rsa.pub pi@sideview1; ssh-copy-id-i~/.ssh/id_rsa.pub pi@sideview2; ssh-copy-id-i ~/.ssh/id_rsa.pub pi@topviewpi)
b. If asked “The authenticity of host ‘remote-hosť can’t be established. Are you sure you want to continue connecting?” enter “yes”.
c. Enter the Raspberry Pi password when prompted (the password is “raspberry” if not altered from default).
4. Open a Terminal window, and enter “cd Desktop” to change directory to the desktop. Create folders on each of the Raspberry Pis to temporarily store images on the Raspberry Pi desktop. To facilitate identification of image source, each folder can be given the respective Raspberry Pi’s hostname (octagon, sideview1, sideview2, topviewpi).
5. For each of the Raspberry Pis, open the piPicture.py script located at Desktop>apps-phenotyping>appendix.4.octagon.multi-image>piPicture.py. Change the picPath directory in line 37 of piPicture.py to the folder created in step 4 (e.g. /home/pi/Desktop/octagon).
6. Similarly, for each of the Raspberry Pis, open the syncScript.sh script located at Desktop>apps-phenotyping>appendix.4.octagon.multi-image>syncScript.sh. Change the rsync local directory in line 1 of syncScript.sh to the folder created in step 4 (e.g. /home/pi/Desktop/octagon).
7. Change the rsync remote directory in line 1 of syncScript.sh to the directory in the remote host where the images will be stored (e.g. jdoe@serverx:/home/jdoe/octagon_images). Make sure that the specified directory exists on the remote host. Remember that passwordless SSH (Appendix 1) must be set up to allow the Raspberry Pi to push data to the remote host.
8. Change ‘<hostname>‘ in line 7 of syncScript.sh to the respective Raspberry Pi hostname.
9. Lines 44 to 58 of piPicture.py script set camera parameters using the Picamera package ((Jones, 2017/2)). These may need to be adjusted depending on the lighting in the octagon chamber, please refer to the Picamera documentation (http://picamera.readthedocs.io/en/release-l.10/recipesl.html) for tips on adjusting parameters.
10. Mount the four Raspberry Pis to the octagon using heavy duty velcro and plug them into a power strip.

### Raspberry Pi multi-image octagon operation protocol

1. Turn all Raspberry Pis on. If desired, put the tarp over the top of the octagon for light diffusion.
2. From your computer (laptop is most convenient), SSH into the octagon Raspberry Pi. Type “ssh pi@octagon” in a Terminal window, and enter the password “raspberry”.
3. To begin imaging, in the Terminal type “bash /home/pi/Desktop/apps-phenotyping/appendix.4.octagon.multi-image/sshScript.sh”
4. When prompted “Please scan barcode or type quit to quit”, type plant id (e.g. speciesx_plantl_treatmentl_repl), or scan a plant id in with a barcode scanner.
5. Place the potted plant into a mounted pot within the octagon (see additional notes)
6. Press enter to acquire images, then wait until prompt “Please scan barcode or type quit to quit” appears again.
7. Repeat steps 6 and 7 to acquire another image, or enter “quit” to quit acquiring images.
8. To shut down sideview1, sideview2, and topviewpi Raspberry Pis, in the Terminal type “bash /home/pi/Desktop/apps-phenotyping/appendix.4.octagon.multi-image/shutdown_a ll_pi.sh”. To shut down the octagon Raspberry Pi, in the terminal window type “sudo halt”.

### Additional Notes

- Affixing a pot to the center of the octagon chamber, with color cards affixed to the outside of the stationary pot (white, black and gray; DGK Color Tools, New York, New York, USA) allows a potted plant to be quickly placed in the same relative position to other images.

## LITERATURE CITED

Al-Tamimi, N., C. Brien, H. Oakey, B. Berger, S. Saade, Y.S. Ho, S.M. Schmöckel, et al. 2016. Salinity tolerance loci revealed in rice using high-throughput non-invasive phenotyping. Nature Communications 7: 13342.

Araus, J.L., and J.E. Cairns. 2014. Field high-throughput phenotyping: the new crop breeding frontier. Trends in Plant Science 19: 52–61.

Bodner, G., M. Alsalem, A. Nakhforoosh, T. Arnold, and D. Leitner. 2017. RGB and spectral root imaging for plant phenotyping and physiological research: experimental setup and imaging protocols. Journal of Visualized Experiments: JoVE 126: e56251.

Chen, D., K. Neumann, S. Friedel, B. Kilian, M. Chen, T. Altmann, and C. Klukas. 2014. Dissecting the phenotypic components of crop plant growth and drought responses based on high-throughput image analysis. The Plant Cell 26: 4636–4655.

Fahlgren N., M. Feldman, M.A. Gehan, M.S. Wilson, C. Shyu, D.W. Bryant, S.T. Hill, et al. 2015. A versatile phenotyping system and analytics platform reveals diverse temporal responses to water availability in *Setaria*. Molecular Plant 8: 1520–1535.

Feldman, M.J., R.E. Paul, D. Banan, J.F. Barrett, J. Sebastian, M.-C. Yee, H. Jiang, et al. 2017. Time dependent genetic analysis links field and controlled environment phenotypes in the model C4 grass Setaria. PLOS Genetics 13: el006841.

Figuière, H., and H.U. Niedermann. 2017. Version 2.5.14 (2017-06-05). gPhoto – opensource digital camera access and remote control. GitHub, San Francisco, California, USA. Website https://github.com/gphoto [accessed 22 August 2017].

Goggin, F.L., A. Lorence, and C.N. Topp. 2015. Applying high-throughput phenotyping to plant-insect interactions: picturing more resistant crops. Current Opinion in Insect Science 9: 69–76.

Granier, C., L. Aguirrezabal, K. Chenu, S.J. Cookson, M. Dauzat, P. Hamard, J.-J. Thioux, et al. 2006. PHENOPSIS, an automated platform for reproducible phenotyping of plant responses to soil water deficit in *Arabidopsis thaliana* permitted the identification of an accession with low sensitivity to soil water deficit. The New Phytologist 169: 623–635.

Honsdorf, N., T.J. March, B. Berger, M. Tester, and K. Pillen. 2014. High-throughput phenotyping to detect drought tolerance QTL in wild barley introgression lines. PLOS ONE 9: e97047.

Huang, H., C.Y. Yoo, R. Bindbeutel, J. Goldsworthy, A. Tielking, S. Alvarez, M.J. Naldrett, et al. 2016. PCH1 integrates circadian and light-signaling pathways to control photoperiod-responsive growth in *Arabidopsis*. eLife 5: el3292.

Iyer-Pascuzzi, A.S., O. Symonova, Y. Mileyko, Y. Hao, H. Belcher, J. Harer, J.S. Weitz, and P.N. Benfey. 2010. Imaging and analysis platform for automatic phenotyping and trait ranking of plant root systems. Plant Physiology 152: 1148–1157.

Jahnke, S., J. Roussel, T. Hombach, J. Kochs, A. Fischbach, G. Huber, and H. Scharr. 2016. phenoSeeder - A robot system for automated handling and phenotyping of individual seeds. Plant Physiology 172: 1358–1370.

Jones, D. 2017. Version 1.13 (2017-02-25). Picamera. GitHub, San Francisco, California, USA. Website https://github.com/waveform80/picamera [accessed 01 September 2017].

Leister, D., C. Varotto, P. Pesaresi, A. Niwergall, and F. Salamini. 1999. Large-scale evaluation of plant growth in *Arabidopsis thaliana* by non-invasive image analysis. Plant Physiology and Biochemistry 37: 671–678.

Minervini, M., M.V. Giuffrida, P. Perata, and S.A. Tsaftaris. 2017. Phenotiki: An open software and hardware platform for affordable and easy image-based phenotyping of rosette-shaped plants. The Plant Journal: for Cell and Molecular Biology 90: 204–216.

Monk, S. 2016. Raspberry Pi cookbook: software and hardware problems and solutions. O’Reilly Media, Sebastopol, California, USA.

Mutka, A.M., S.J. Fentress, J.W. Sher, J.C. Berry, C. Pretz, D.A. Nusinow, and R. Bart. 2016. Quantitative, image-based phenotyping methods provide insight into spatial and temporal dimensions of plant disease. Plant Physiology 172: 650–660.

Pauli, D., P. Andrade-Sanchez, A.E. Carmo-Silva, E. Gazave, A.N. French, J. Heun, D.J. Hunsaker, et al. 2016. Field-based high-throughput plant phenotyping reveals the temporal patterns of quantitative trait loci associated with stress-responsive traits in cotton. G3 6: 865–879.

Poorter, H., F. Fiorani, R. Pieruschka, T. Wojciechowski, W.H. van der Putten, M. Kleyer, U. Schurr, and J. Postma. 2016. Pampered inside, pestered outside? Differences and similarities between plants growing in controlled conditions and in the field. The New Phytologist 212: 838–855.

Shafiekhani, A., S. Kadam, F.B. Fritschi, and G.N. DeSouza. 2017. Vinobot and Vinoculer: two robotic platforms for high-throughput field phenotyping. Sensors 17: 214.

Upton, E., and G. Halfacree. 2014. Raspberry Pi user guide. John Wiley & Sons, Hoboken, New Jersey, USA.

Yang, W., Z. Guo, C. Huang, L. Duan, G. Chen, N. Jiang, W. Fang, et al. 2014. Combining high-throughput phenotyping and genome-wide association studies to reveal natural genetic variation in rice. Nature Communications 5: 5087.

Zhang, X., C. Huang, D. Wu, F. Qiao, W. Li, L. Duan, K. Wang, et al. 2017. High-throughput phenotyping and QTL mapping reveals the genetic architecture of maize plant growth. Plant Physiology 173: 1554–1564.

